# Reproductive tradeoffs govern sexually dimorphic tubular lysosome induction in *C. elegans*

**DOI:** 10.1101/2022.03.16.484652

**Authors:** Cara D. Ramos, K. Adam Bohnert, Alyssa E. Johnson

## Abstract

Animals of different sexes often exhibit unique behaviors that benefit their specific reproductive interests. In the nematode *Caenorhabditis elegans*, self-fertilizing hermaphrodites can reproduce without a mate and thus prioritize feeding to satisfy the high energetic costs of reproduction. However, males, which rely on finding potential mates for reproduction, sacrifice feeding and instead prioritize exploratory behavior. Here, we demonstrate that these differences in behavior are linked to sexual dimorphism at the cellular level; young males raised on a rich food source show constitutive induction of gut tubular lysosomes, a non-canonical lysosome morphology that typically forms in the gut of young hermaphrodites only when food is limited. We find that male-specific induction of gut tubular lysosomes on abundant food is due to self-imposed dietary restriction through *daf-7*/TGFβ signaling, which promotes mate-searching at the cost of feeding. While gut tubular lysosomes are largely absent from well-fed hermaphrodites at the start of adulthood, their induction accelerates in hermaphrodites in early aging, dependent on the presence of sperm and, partly, on embryo production. These findings identify tubular lysosome induction as a sexually dimorphic cellular event that may integrate animal physiology with sex-specific behavioral differences important for reproductive success.

## Introduction

Many animal traits and behaviors, especially those linked to reproduction, display sexual dimorphism (Portman, 2007; Yamamoto, 2007; Zilkha et al., 2021). Such phenotypes vary from one sex to another within a single species, and maintenance of these differences is often vital for efficient reproduction and, ultimately, species survival. In the nematode *C. elegans*, the choice between feeding and mate searching presents an interesting example of a sexually dimorphic behavior; young male worms prioritize mate searching over feeding, whereas hermaphrodites, which reproduce on their own using self-sperm and oocytes, constantly prioritize feeding (Lipton et al., 2004; Ryan et al., 2014). The male-specific preference for mating over feeding is controlled by the *daf-7/*TGFβ neuroendocrine signaling axis. In well-fed young males, elevated DAF-7 inhibits expression of the odorant receptor *odr-10*, thereby reducing the preference to feed (Hilbert and Kim, 2017; Wexler et al., 2020). In contrast, inhibiting *daf-7* in young males is sufficient to prevent mate-searching behavior and to promote feeding behavior instead (Wexler et al., 2020). While sacrificing feeding for exploratory behavior is a critical element of male reproductive behavior and success, its effects on other aspects of animal physiology is less clear. In principle, metabolic parameters linked to nutritional status could be impacted in males. To what degree this occurs, and how it compares to changes in hermaphrodites, which assume high metabolic costs in producing embryos, is unknown.

In previous work, we demonstrated that nutritional cues govern the induction of autophagic tubular lysosomes (TLs) in the digestive tissues of worms and flies (Dolese et al., 2021; Villalobos et al., 2021). Upon starvation or dietary restriction (DR), gut lysosomes transform from vesicles into expansive tubular networks that show high degradative activity (Dolese et al., 2021; Villalobos et al., 2021). Importantly, this morphological transformation in lysosome structure supports lifespan extension in nutrient deprived conditions, and can even be artificially mimicked in well-fed animals for health benefits (Villalobos et al., 2021). Given that nutritional cues are intimately linked to sexually dimorphic feeding/mating behaviors in *C. elegans*, it is conceivable that TL induction might naturally vary between the biological sexes in this species. If so, this could contribute to sex-specific differences in animal health and physiology.

Here, we demonstrate that young male *C. elegans* induce TLs in their gut even in the presence of abundant nutrient sources. We further demonstrate that this is linked to the self-imposed dietary avoidance that permits male worms to spend more time searching for a mate. In contrast, hermaphrodites show lower TL-related signaling in young adulthood, but this increases dramatically, surpassing even the male, during aging. Using sperm-defective mutants, we find that elevated TL induction with age in hermaphrodites relates to both the presence of sperm and embryo production, potentially as a mechanism to supply nutrients to the developing progeny and/or the mother. Collectively, our results suggest that reproductive tradeoffs dictate TL induction in *C. elegans* and may provide physiological support to animals as they prioritize distinct modes of reproductive success.

## Materials and Methods

### Strains

Table S1 provides a complete list of strains used in this study. Endogenously-tagged *spin-1::mCherry* was generated by *In Vivo* Biosystems using CRISPR technology. For genetic crosses, the endogenous *spin-1::mCherry* transgene was tracked by stereomicroscopy, and genetic mutations were verified by phenotypic characterization and/or sequencing.

### Animal maintenance

Unless otherwise noted, worms were raised at 20ºC on NGM agar (51.3 mM NaCl, 0.25% peptone, 1.7% agar, 1 mM CaCl_2_, 1 mM MgSO_4_, 25 mM KPO_4_, 12.9 µM cholesterol, pH 6.0). For standard experiments, fed worms were maintained on NGM agar plates that had been seeded with *E. coli* OP50 bacteria. To obtain starved adult worms, worms were washed 5x in 5 mL M9 buffer and transferred onto NGM agar that lacked OP50 bacteria. Synchronous populations of worms were obtained by bleaching young-adult hermaphrodites. Briefly, adult hermaphrodites were vortexed in 1 mL bleaching solution (0.5 M NaOH, 20% bleach) for 5 minutes to isolate eggs, and eggs were then washed three times in M9 buffer (22 mM KH_2_PO_4_, 42 mM Na_2_HPO_4_, 85.5 mM NaCl, 1 mM MgSO_4_) before plating.

For RNAi experiments, synchronous populations of animals were grown on OP50-seeded NGM plates until late L4 or day 1 of adulthood, at which time they were transferred to RNAi plates (NGM plus 100 ng/µl carbenicillin and 1 mM IPTG) that had been seeded with bacteria expressing the relevant RNAi clone. An empty L4440 vector was used as a negative control.

In experiments involving the *fog-2* strain, virgin females were isolated by transferring feminized individuals onto NGM plates seeded with OP50 bacteria without males. Populations of mated feminized worms were kept on plates with young males throughout the experiments.

Sperm-defective *fer-1* and *spe-9* strains are fertile at 15ºC but, when raised at 25ºC, produce sperm that signal appropriately but are defective in fertilizing oocytes. In experiments involving these mutants, strains were routinely maintained at 15ºC until the experiment was conducted. To obtain experimental, synchronous populations of *fer-1* and *spe-9* mutants, strains were bleached, and NGM agar plates with eggs and OP50 bacteria were shifted to 25ºC to render animals self-sterile. Control strains for these experiments were treated identically at the same time.

For aging experiments, synchronous populations of worms were obtained by bleaching young adult hermaphrodites and plating their eggs onto NGM plates seeded with OP50 bacteria. For progeny-producing strains, the sample populations were transferred to fresh NGM plates every two days to isolate them from their progeny and to ensure a continuous food supply.

### Male generation and propagation

Males were generated by heat shocking hermaphrodites. Briefly, hermaphrodite animals were subjected to a persistent heat shock at 25ºC, or, alternatively, L4 hermaphrodites were subjected to a briefer 4-6 hour heat shock at 30ºC. Male progeny isolated in the next generation were propagated by mating. For mating, 8-10 hermaphrodites were placed on a 35 mm NGM plate with roughly 20 males and maintained at 20ºC overnight. This plate was seeded with a small scoop of OP50 bacteria at the center of the plate to increase the likelihood of mating encounters. On the following day, hermaphrodites were transferred to 60 mm NGM plates and allowed to lay eggs. The mating process could be repeated iteratively in subsequent generations to maintain a consistent population of males.

### Male-conditioned plates

30 male worms were transferred onto NGM plates seeded with OP50 bacteria to allow males to secrete pheromones onto the plates (Maures et al., 2014). After two days, males were transferred off the plates, and *fog-2* feminized virgins were transferred onto the plates at day one of adulthood to expose them to the male-conditioned environment. *fog-2* feminized virgins were imaged two or four days after exposure.

### Microscopy

4% agarose (Fisher Bioreagents) pads were dried on a Kimwipe (Kimtech) and then placed on top of a Gold Seal™ glass microscope slide (ThermoFisher Scientific). A small volume of 20 mM levamisole (Acros Organics) was spotted on the agarose pad. Worms were transferred to the levamisole spot, and a glass cover slip (Fisher Scientific) was placed on top to complete the mounting. Live-animal fluorescence microscopy was performed using a Leica DMi8 THUNDER imager, equipped with 10X (NA 0.32), 40X (NA 1.30), and 100X (NA 1.40) objectives and GFP and Texas Red filter sets.

### Image analysis

Images were processed using LAS X software (Leica) and FIJI/ImageJ (NIH). Lysosome networks were analyzed using “Skeleton” analysis plugins in FIJI. Briefly, images were converted to binary 8-bit images and then to skeleton images using the “Skeletonize” plugin. Skeleton images were then quantified using the “Analyze Skeleton” plugin. Number of objects, number of junctions, and object lengths were scored. An “object” is defined by the Analyze Skeleton plugin as a branch connecting two endpoints, an endpoint and a junction, or two junctions. Junctions/object was used as a parameter to quantify network integrity. For SPIN-1:mCherry fluorescence quantification, the gut tissue was outlined using the free-draw tool in FIJI/ImageJ, and average fluorescence intensity of the outlined area was measured. For all fluorescence intensity experiments, the same laser intensity (50%), exposure time (300 ms), and FIM (100%) were used.

### Statistical analyses

Data were statistically analyzed using GraphPad Prism. For two sample comparisons, an unpaired t-test was used to determine significance (α=0.05). For three or more samples, a one-way ANOVA with Dunnett’s multiple comparisons was used to determine significance (α=0.05).

## Results and Discussion

### Young male worms show constitutive TL induction in the gut, even under nutrient rich conditions

Previously, we demonstrated that well-fed *C. elegans* hermaphrodites show gut lysosomes that are morphologically static and predominantly vesicular in structure; however, upon starvation, these lysosomes transform into dynamic, autophagic, tubular networks (Dolese et al., 2021; Villalobos et al., 2021). Thus, starvation acts as a natural trigger for TL induction in the gut of *C. elegans* hermaphrodites. To extend these studies, we explored whether male animals, which normally sacrifice feeding for mating, show differences in TL induction, perhaps even in the presence of food. We tracked lysosomes in males on and off of food using endogenous *spin-1::mCherry*, which encodes a Spinster ortholog that robustly labels TLs (Villalobos et al., 2021). We found that young male worms, unlike young hermaphrodites (Villalobos et al., 2021), in fact exhibited TLs in the gut when food was abundant (Figure 1A-B). As in starved hermaphrodites (Villalobos et al., 2021), TL induction in young males on food was accompanied by a relative increase in endogenous SPIN-1 protein intensity (Figure 1C-D), suggesting SPIN-1 protein expression serves as a proxy for TL induction. Additionally, young male worms that were raised without food exhibited no further increase in SPIN-1::mCherry protein intensity compared to males raised on food (Figure 1E-F). Thus, in young males, TLs appear to be constitutively induced in the gut, regardless of food status.

**Figure 1:**
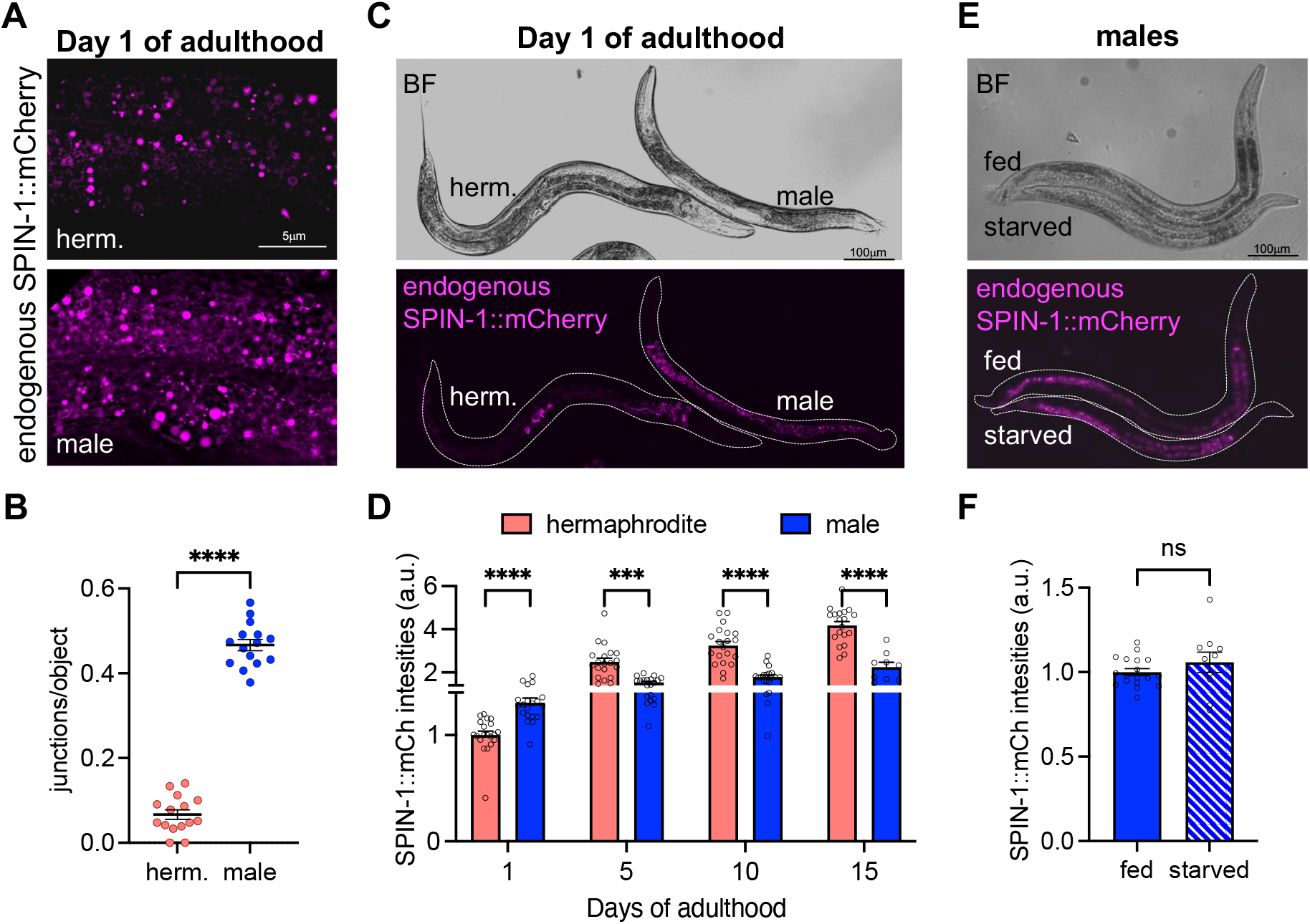
Young male worms show constitutive TL induction in the gut, even under nutrient rich conditions. **A**. Representative images of endogenously tagged SPIN-1::mCherry in hermaphrodite and male worms. **B**. Quantification of lysosome junctions/object in hermaphrodite and male worms. **C**. Representative images of *spin-1* expression in hermaphrodite and male worms. **D**. Quantification of SPIN-1::mCherry intensities in hermaphrodite and male worms throughout adulthood. **E**. Representative images of *spin-1* expression in fed and starved male worms. **F**. Quantification of SPIN-1::mCherry intensities in fed and starved male worms.

Like starvation, aging also induces gut TLs in hermaphrodite worms (Dolese et al., 2021; Villalobos et al., 2021). Given that young male worms exhibited TL induction and higher SPIN-1::mCherry fluorescence intensity compared to young hermaphrodites, we surmised that male worms might have a higher basal level of SPIN-1, which would continue to increase relative to hermaphrodite levels during aging. However, this was not the case; by day five of adulthood, SPIN-1::mCherry intensity in hermaphrodites superseded that in males, and this trend continued into late life (Figure 1D). Thus, the stronger TL induction in nutrient rich conditions was specific to young male worms. Moreover, some biological activities specific to hermaphrodites in young adulthood may contribute to their relatively fast increase in SPIN-1 protein levels with age.

### *Elevated TL induction in young males results from* daf-7*-dependent prioritization of mating over feeding*

Our observation that TLs were induced in male worms even on a rich food source could suggest that the same starvation-based mechanisms of TL induction seen in hermaphrodites do not apply to the male sex. Yet, given the consideration that young male worms trade off feeding in order to spend more time searching for a mate (Lipton et al., 2004; Ryan et al., 2014), we reasoned that a self-imposed DR due to prioritization of exploratory behavior may explain the constitutive TL induction in males raised on food. To test this hypothesis, we manipulated the *daf-7/*TGFβ signaling axis that differentially regulates feeding/mating decision-making in *C. elegans* hermaphrodites and males (Figure 2A) (Hilbert and Kim, 2017; Milward et al., 2011; Wexler et al., 2020; You et al., 2008). Strikingly, inhibition of *daf-7* by RNAi prevented the male-specific increase in SPIN-1::mCherry fluorescence intensities and TL induction during young age (Figure 2B-C). These data support the notion that TL induction in young male worms is a consequence of a self-imposed DR caused by prioritized mate-searching behaviors.

**Figure 2:**
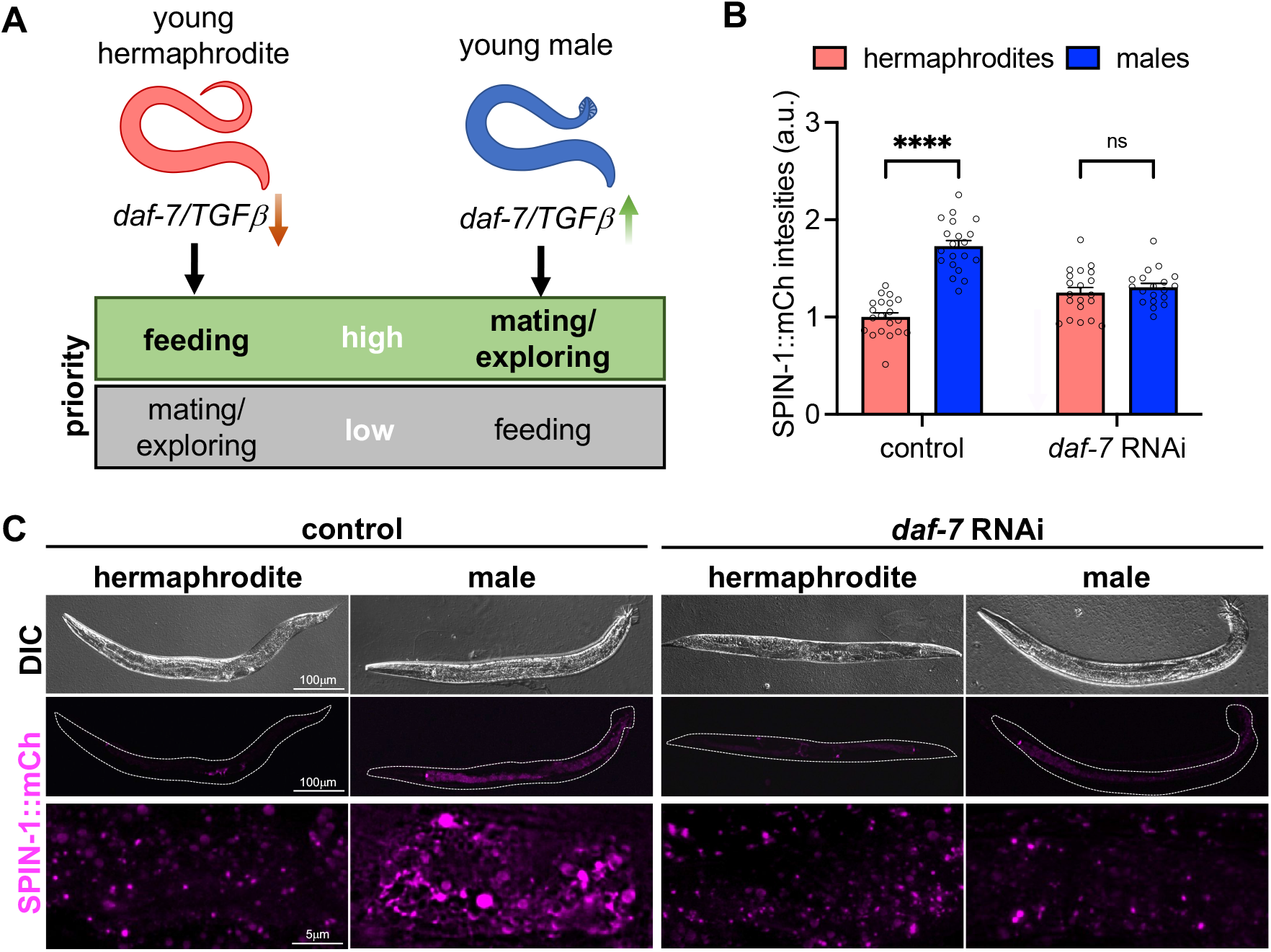
Elevated TL induction in young males results from *daf-7*-dependent prioritization of mating over feeding. **A**. Schematic diagram illustrating the *daf-7*-dependent sexually dimorphic feeding/exploring behavior in *C. elegans*. In young hermaphrodites, *daf-7* signaling is downregulated, which promotes feeding behaviors. In contrast, *daf-7* signaling is upregulated in young male worms to promote exploratory behaviors. **B**. Quantification of SPIN-1::mCherry intensities in control and *daf-7* RNAi treated worms. **C**. Representative images of *spin-1* expression and TL induction in control and *daf-7* RNAi treated worms.

### Sperm signaling and embryo production contribute to increased SPIN-1 intensities in mothers during early aging

We next considered mechanisms that possibly contribute to elevated SPIN-1 protein expression and TL induction in hermaphrodites with age. Although the two natural sexes of *C. elegans* are hermaphrodite (XX) and male (XO), “feminized” worms can be obtained using XX animals incapable of producing sperm (Barton and Kimble, 1990). For example, in *fog-2* mutant animals, germ cells that would normally differentiate into sperm instead differentiate into oocytes (Schedl and Kimble, 1988). Using SPIN-1::mCherry intensity levels as a proxy for TL induction, we compared SPIN-1::mCherry intensities in hermaphrodites, virgin feminized animals, and mated feminized animals throughout adulthood. At day one, no significant differences were observed between the three groups (Figure 3A-B). However, by days five and ten, SPIN-1::mCherry intensities were significantly lower in virgin feminized animals compared to both hermaphrodite and mated feminized animals (Figure 3A-B). These data suggest that the presence of sperm might drive the steep increase in hermaphrodite SPIN-1 expression during adulthood.

**Figure 3:**
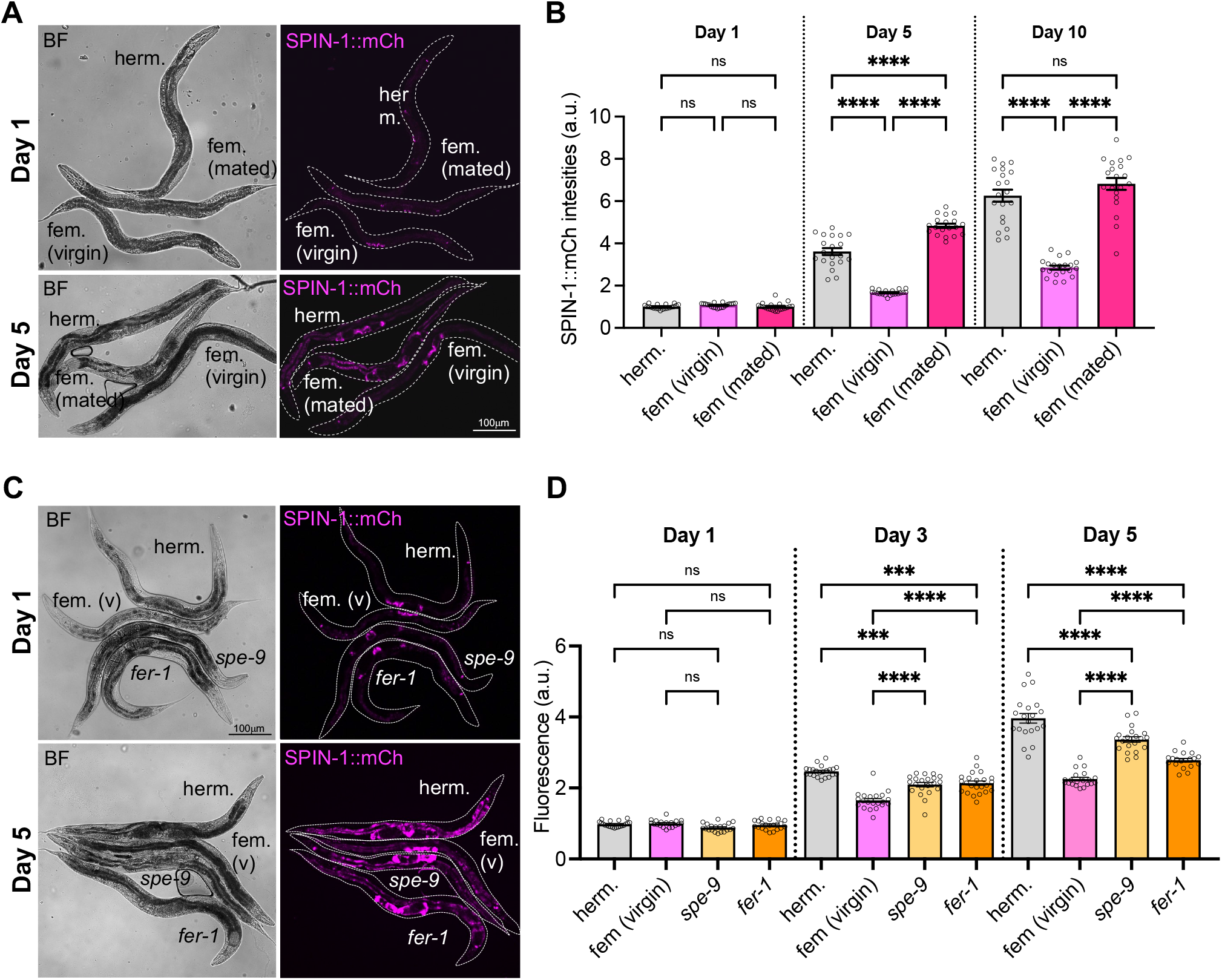
Sperm signaling and embryo production contribute to increased SPIN-1 intensities in mothers during early aging. **A-B**. Representative images of *spin-1* expression (A) and quantification of SPIN-1::mCherry intensities (B) in hermaphrodite and feminized worms (virgin and mated) throughout adulthood. **C-D**. Representative images of *spin-1* expression (C) and quantification of SPIN-1::mCherry intensities (D) in hermaphrodites, virgin feminized worms, and mutants with fertilization-incompetent sperm (*spe-9* and *fer-1*) during early adulthood.

Intriguingly, the mere presence of mating-competent male worms has been demonstrated to depreciate physiological health and lifespan of hermaphrodite worms cultured in the same environment (Maures et al., 2014). Moreover, pre-conditioning plates with male pheromones alone is sufficient to cause reduced lifespan in hermaphrodites, indicating that exposure to male pheromones rather than mating triggers accelerated aging phenotypes in hermaphrodite worms (Maures et al., 2014). These studies prompted us to test whether exposure to male pheromones was also sufficient to induce age-related changes to *spin-1* expression levels in feminized animals. We found that virgin feminized worms exposed to the male-conditioned plates failed to exhibit increased SPIN-1::mCherry intensities compared to control virgin feminized worms after either two or four days of exposure to male-conditioned plates (Figure S1A-B). Thus, exposure to male-specific pheromones is insufficient to induce SPIN-1 levels, consistent with sperm instead playing a causal role.

The above results suggested three possibilities: (i) signals emanating from sperm trigger an age-related increase in SPIN-1 and TLs in the mother, independent of fertilization; (ii) production of embryos upon fertilization of oocytes by sperm instead triggers TL induction; or (iii) signals from both sperm and embryo production contribute to increased SPIN-1 and TL induction. To distinguish between these possibilities, we examined SPIN-1::mCherry intensities in *spe-9* and *fer-1* mutants, which can produce both gametes (sperm and oocytes) but have mutations that render the sperm incapable of fertilization (L’Hernault et al., 1988; Singson et al., 1998; Ward and Miwa, 1978; Ward et al., 1981). These sperm-defective mutants allowed us to examine a biological scenario in which sperm signals are present, but embryo production is disabled. At day one of adulthood, no significant increase in SPIN-1::mCherry intensity was detected in either *spe-9* and *fer-1* mutants compared to virgin feminized worms (Figure 3C-D). However, by days three and five, SPIN-1::mCherry intensity in *spe-9* and *fer-1* mutants increased significantly compared to virgin feminized worms, albeit not to the level of hermaphrodite worms (Figure 3C-D). These results suggest that signals from both sperm and embryo production contribute to increasing SPIN-1::mCherry levels and TL induction in the mother.

In conclusion, we have uncovered two sexually dimorphic properties of TL induction in *C. elegans*: (1) young males show constitutive TL induction due to a self-imposed DR that permits enhanced mate-searching behavior; and (2) TL induction in hermaphrodites commences later during aging, dependent on previous reproductive signaling and activity. We propose that TLs are induced by different mechanisms in each sex to meet the nutritional demands imposed by their distinct reproductive activities. In young males, the induction of TLs could provide health benefits during this self-imposed DR period to boost their reproductive fitness. DR has been long known to confer health benefits and extend lifespan in many species; however, in *C. elegans*, lifespan is extended by DR in hermaphrodites, but not in males (Honjoh et al., 2017). This supports the notion that male worms, which are calorically restricted by choice, already exhibit the health benefits of DR as a natural consequence of this behavior and, thus, do not exhibit any further lifespan extension when put under experimental dietary constraints. Moreover, we have shown previously that artificial induction of TLs allows worms to sustain their mobility longer in life (Villalobos et al., 2021). Thus, it is interesting to speculate whether induction of TLs in young males improves their physical fitness and ability to find a mate. In the case of hermaphrodites, developing embryos inside the uterus require significant nutritional support, which must come from the mother. Thus, TL induction might allow mothers to recycle nutrients, such that they can provide additional nutritional sustenance to the developing embryos and/or themselves during reproduction. Collectively, these findings add to growing evidence indicating that different sexes have distinct nutritional requirements during their reproductive lifespan, and they also suggest that TL induction may contribute to sustaining reproductive fitness by different mechanisms in each sex.

## Acknowledgments

The authors thank all members of the Bohnert and Johnson labs for helpful discussions on this project.

## Competing interests

The authors declare that they have no competing interests.

## Author contributions

Conceptualization: KAB, AEJ; Methodology: CDR; Investigation: CDR, KAB, AEJ; Visualization: CDR, KAB, AEJ; Funding acquisition: KAB, AEJ; Project administration: KAB, AEJ; Supervision: KAB, AEJ; Writing – original draft: KAB, AEJ; Writing – review &editing: CDR, KAB, AEJ

## Funding

LSU Office of Research and Economic Development, the LSU College of Science, and the LSU Department of Biological Sciences (KAB, AEJ); the W.M. Keck Foundation (KAB, AEJ) and the National Institutes of Health (AEJ; R35GM138116).

## Notes

### Competing Interest Statement

The authors have declared no competing interest.

## References

Akagi, K., Wilson, K.A., Katewa, S.D., Ortega, M., Simons, J., Hilsabeck, T.A., Kapuria, S., Sharma, A., Jasper, H., and Kapahi, P. (2018). Dietary restriction improves intestinal cellular fitness to enhance gut barrier function and lifespan in D. melanogaster. PLOS Genet. 14, e1007777.

Barton, M.K., and Kimble, J. (1990). fog-1, a regulatory gene required for specification of spermatogenesis in the germ line of Caenorhabditis elegans. Genetics 125, 29–39.

Dolese, D.A., Junot, M.P., Ghosh, B., Butsch, T.J., Johnson, A.E., and Bohnert, K.A. (2021). Degradative tubular lysosomes link pexophagy to starvation and early aging in C. elegans. Autophagy.

Hilbert, Z.A., and Kim, D.H. (2017). Sexually dimorphic control of gene expression in sensory neurons regulates decision-making behavior in C. elegans. Elife 6.

Honjoh, S., Ihara, A., Kajiwara, Y., Yamamoto, T., and Nishida, E. (2017). The Sexual Dimorphism of Dietary Restriction Responsiveness in Caenorhabditis elegans. Cell Rep. 21, 3646–3652.

Kapahi, P., Kaeberlein, M., and Hansen, M. (2017). Dietary restriction and lifespan: Lessons from invertebrate models. Ageing Res. Rev. 39, 3–14.

L’Hernault, S.W., Shakes, D.C., and Ward, S. (1988). Developmental genetics of chromosome I spermatogenesis-defective mutants in the nematode Caenorhabditis elegans. Genetics 120, 435–452.

Lakowski, B., and Hekimi, S. (1998). The genetics of caloric restriction in Caenorhabditis elegans. Proc. Natl. Acad. Sci. U. S. A. 95, 13091.

Lipton, J., Kleemann, G., Ghosh, R., Lints, R., and Emmons, S.W. (2004). Mate searching in Caenorhabditis elegans: a genetic model for sex drive in a simple invertebrate. J. Neurosci. 24, 7427–7434.

Magwere, T., Chapman, T., and Partridge, L. (2004). Sex differences in the effect of dietary restriction on life span and mortality rates in female and male Drosophila melanogaster. J. Gerontol. A. Biol. Sci. Med. Sci. 59, 3–9.

Maures, T.J., Booth, L.N., Benayoun, B.A., Izrayelit, Y., Schroeder, F.C., and Brunet, A. (2014). Males shorten the life span of C. elegans hermaphrodites via secreted compounds. Science 343, 541–544.

Milward, K., Busch, K.E., Murphy, R.J., De Bono, M., and Olofsson, B. (2011). Neuronal and molecular substrates for optimal foraging in Caenorhabditis elegans. Proc. Natl. Acad. Sci. U. S. A. 108, 20672–20677.

Portman, D.S. (2007). Genetic Control of Sex Differences in C. elegans Neurobiology and Behavior. Adv. Genet. 59, 1–37.

Ryan, D.A., Miller, R.M., Lee, K., Neal, S.J., Fagan, K.A., Sengupta, P., and Portman, D.S. (2014). Sex, age, and hunger regulate behavioral prioritization through dynamic modulation of chemoreceptor expression. Curr. Biol. 24, 2509–2517.

Schedl, T., and Kimble, J. (1988). fog-2, a germ-line-specific sex determination gene required for hermaphrodite spermatogenesis in Caenorhabditis elegans. Genetics 119, 43–61.

Singson, A., Mercer, K.B., and L’Hernault, S.W. (1998). The C. elegans spe-9 Gene Encodes a Sperm Transmembrane Protein that Contains EGF-like Repeats and Is Required for Fertilization. Cell 93, 71–79.

Villalobos, T. V., Ghosh, B., Alam, S., Butsch, T.J., Mercola, B.M., Ramos, C.D., Das, S., Eymard, E.D., Bohnert, K.A., and Johnson, A.E. (2021). Tubular lysosome induction couples animal starvation to healthy aging. BioRxiv 2021.10.28.466256.

Ward, S., and Miwa, J. (1978). Characterization of temperature-sensitive, fertilization-defective mutants of the nematode caenorhabditis elegans. Genetics 88, 285–303.

Ward, S., Argon, Y., and Nelson, G.A. (1981). Sperm morphogenesis in wild-type and fertilization-defective mutants of Caenorhabditis elegans. J. Cell Biol. 91, 26–44.

Weindruch, R., Walford, R.L., Fligiel, S., and Guthrie, D. (1986). The retardation of aging in mice by dietary restriction: longevity, cancer, immunity and lifetime energy intake. J. Nutr. 116, 641–654.

Wexler, L.R., Miller, R.M., and Portman, D.S. (2020). C. elegans Males Integrate Food Signals and Biological Sex to Modulate State-Dependent Chemosensation and Behavioral Prioritization. Curr. Biol. 30, 2695–2706.e4.

Yamamoto, D. (2007). The Neural and Genetic Substrates of Sexual Behavior in Drosophila. Adv. Genet. 59, 39–66.

You, Y. jai, Kim, J., Raizen, D.M., and Avery, L. (2008). Insulin, cGMP, and TGF-β Signals Regulate Food Intake and Quiescence in C. elegans: A Model for Satiety. Cell Metab. 7, 249–257.

Zilkha, N., Sofer, Y., Kashash, Y., and Kimchi, T. (2021). The social network: Neural control of sex differences in reproductive behaviors, motivation, and response to social isolation. Curr. Opin. Neurobiol. 68, 137–151.

